# Electrochemical disruption of extracellular electron transfer inhibits *Pseudomonas aeruginosa* cell survival within biofilms and is synergistic with antibiotic treatment

**DOI:** 10.1101/2022.09.15.508205

**Authors:** Fernanda Jiménez Otero, Dianne K. Newman, Leonard M. Tender

## Abstract

Survival of cells within oxygen-limited regions in *Pseudomonas aeruginosa* biofilms is enabled by using small redox active molecules as electron shuttles to access distal oxidants. This respiratory versatility makes *P. aeruginosa* biofilms common in chronic wound infections and recalcitrant to treatment. Here, we show that electrochemically controlling the redox state of these electron shuttles, specifically pyocyanin, can impact cell survival within anaerobic *P. aeruginosa* biofilms and can act synergistically with antibiotic treatment. We inhibited pyocyanin redox cycling under anoxic conditions by blocking its ability to be re-oxidized and thus serve as an electron shuttle via poising an electrode at a reductive potential that cannot regenerate oxidized pyocyanin (*i.e*. −400mV vs Ag/AgCl). This resulted in a decrease in CFUs within the biofilm of 100x compared to samples exposed to an electrode poised at an oxidizing potential that permits pyocyanin re-oxidation (*i.e.* +100mV vs Ag/AgCl). Phenazine-deficient Δ*phz\** biofilms were not affected by the redox potential of the electrode, but were re-sensitized by adding pyocyanin. The effect of EET disruption was exacerbated when biofilms were treated with sub-MICs of a range of antibiotics. Most notably, 4 μg/ml of the aminoglycoside gentamicin in a reductive environment almost completely eradicated wild type biofilms but had no effect on the survival of Δ*phz*\* biofilms, suggesting reduced phenazines are toxic, and combined with antibiotic treatment can lead to extensive killing.

**Importance:** Biofilms provide a protective environment but they also present challenges to the cells living within them, such as overcoming diffusion limitation of nutrients and oxygen. *Pseudomonas aeruginosa* overcomes oxygen limitation by secreting soluble redox active molecules as electron shuttles to access distal oxygen. Here, we show that electrochemically blocking the redox cycling of one of these electron shuttles, pyocyanin, decreases cell survival within biofilms and acts synergistically with gentamicin to kill cells. Our results highlight the importance of the role that the redox cycling of electron shuttles fulfills within *P. aeruginosa* biofilms.

## Observation

Biofilms provide bacterial cells with a protective environment where persistence and antibiotic tolerance arise, making them a leading contributor to chronic infections (1). Extracellular electron transfer (EET) pathways have been recurrently found among biofilm-forming opportunistic pathogens (2–4). Such pathways are often dependent on the redox cycling of either self-made or borrowed small molecules that serve as electron shuttles to extracellular terminal electron acceptors (5). Specifically, in the biofilms formed by *Pseudomonas aeruginosa* PA14 (6), oxygen limitation within anoxic regions is overcome through the use of phenazines as electron shuttles to reduce distal oxygen (7, 8). Of the different phenazines produced by *P. aeruginosa* PA14, pyocyanin (PYO) is present at high abundance and facilitates EET via its association with extracellular DNA (9).

Electrochemical control over biofilms has been explored in several ways in the past. Applying a weak current to biofilms formed on electrodes by persister *P. aeruginosa* PAO1 cells decreases cell survival, but the mechanism underpinning this observation is not understood (10). Electric bandages poised at - 600 mV vs Ag/AgCl to produce H2O2 also decrease cell survival within multi-species biofilms (11). However, H2O2 has been known to prolong the wound healing process and is cytotoxic at high concentrations, which may be counteractive to its antimicrobial effects (12). Providing a poised electrode as alternative electron acceptor in the proximity of agar-grown *P. aeruginosa* PA14 colonies delays wrinkling colony morphology associated with the development of oxygen-limited regions by alleviating oxidant limitation (13). Additionally, biochemically altering PYO through demethylation has also been effective in decreasing *P. aeruginosa* PA14 cell survival and is synergistic with antibiotic treatment (14).

Here, we report that disrupting extracellular electron transfer electrochemically under anoxic conditions can inhibit cell survival. We hypothesized that, in the absence of oxygen, blocking redox cycling of electron shuttles during extracellular electron transfer would negatively affect cell survival within *P. aeruginosa* PA14 biofilms. Biofilms were grown for 5 days in actively aerated 3-electrode electrochemical reactors using an indium tin oxide (ITO)-covered glass slide as both biofilm attachment surface and working electrode. In contrast to previous studies (15), under our conditions, the only phenazine detected was pyocyanin (Fig. S1 and Detailed Experimental Procedures). Electrode-attached biofilms were then transferred to anoxic reactors for 72 hours, after which biofilms were harvested for colony forming unit (CFU) counts and biofilm imaging (Fig 1A). Throughout each experiment, the ITO working electrode was poised at either the PYO-oxidative potential of +100 mV vs Ag/AgCl, or the PYO-reductive potential of −400 mV vs Ag/AgCl, which is not low enough to produce H2O2 (16). Under PYO-oxidative conditions, electron shuttling occurs between cells and the electrode serving as terminal electron acceptor. Under PYO-reductive conditions, electron shuttling does not occur because oxidized PYO cannot be regenerated (Fig. 1B). “Untreated” control biofilms were set to open circuit (OC) conditions, which only act as redox potential-monitoring assays of the electrode/biofilm interface and for which electron shuttling only occurs with trace oxygen (present in all conditions but its effect is expected to be negligible under oxidative conditions and outcompeted by the electrode under reductive conditions).

**Figure 1.**
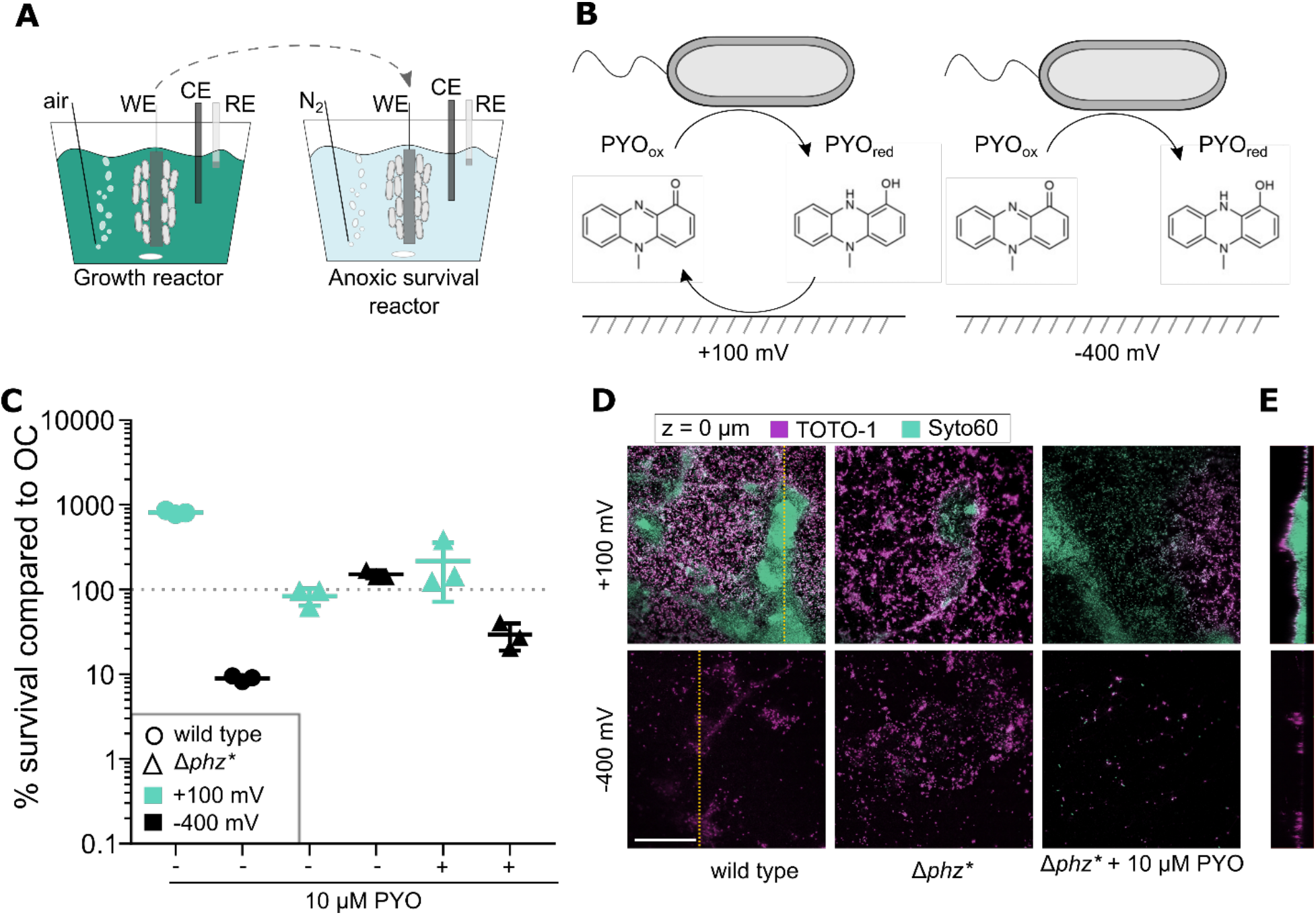
EET impacts cell survival and biofilm morphology. **A**) Schematic representation of experimental setup. Biofilm attachment surface served as the working electrode and was poised at either open circuit (OC), PYO-oxidative potential (+100 mV, or PYO-reductive potential (−400 mV vs Ag/AgCl). Growth reactor was incubated under oxic conditions for 5 days with fresh medium exchanged every 24 hours. The working electrode was then transferred to an anoxic survival reactor flushed with N_2_ gas and incubated for 72 hours before harvesting and processing biofilm. Abbs: WE=working electrode, CE=counter electrode, RE=reference electrode. **B**) Schematic representation of redox cycling of PYO between cells and electrode under PYO-oxidative (+100 mV) and PYO-reductive (−400 mV) conditions; cycling occurs in the former but not the latter condition. **C**) CFUs after 72 hours under anoxic conditions (n =3) normalized to parallel OC negative control samples for wild type, Δ*phz\**, and Δ*phz\** + 10 μM PYO biofilms. Error bars represent standard error. **D**) Fluorescence microscopy images of biofilm interface with electrode surface using TOTO-1 (cell-impermeable, eDNA) and Syto60 (cell-permeable, all DNA) from samples representative of 3 fields of view from triplicate cultures. Bar = 50 um. **E**) Vertical slice of complete z-stack along yellow dotted lines shown in wild type panels in D).

### Electrochemically blocking extracellular electron transfer during anoxic conditions decreases cell survival

Compared to OC control conditions, anoxic PYO-oxidative conditions enhanced cell survival by ~10x while PYO-reductive conditions decreased wild type cell survival by ~10x (CFUs/cm^2^ ± standard error (SE), n = 3, for OC = (2.35 ± 0.11) x 10^4^, +100 mV = (1.91 ± 0.06) x 10^5^, and −400 mV = (2.10 ± 0.09) x 10^3^, Fig 1C). These results support our hypothesis that the electrode can support EET and enhance cell survival when poised as an anode (+100 mV) and can cause decrease in cell survival when poised as a cathode (−400 mV) by either 1) blocking EET and outcompeting PYO oxidation by trace oxygen, or 2) increasing the proportion of reduced PYO, which may be toxic to cells. This trend held for biofilms grown on glass surfaces within PYO-reductive/oxidative reactors placed ~3 cm from the working electrode (Fig. S2), suggesting that electron shuttling enabled by an electrode at a distance can be effective, as expected from previous studies on liquid cultures (7). Phenazine-deficient Δ*phz\** biofilms were not affected by the redox potential of the biofilm surface but were sensitized under PYO-reductive conditions by the addition 10 μM PYO, indicating that cell survival in this context is PYO-mediated EET-dependent (Fig. 1 and S1).

Biofilm morphology was qualitatively consistent with results from CFU counts, with biofilms treated under PYO-oxidative conditions showing full electrode surface coverage and secondary structures up to 100 μm thick; large micro colonies stained brightly with SYTO 60 in the core yet took up TOTO-1 in the periphery (Fig. 1D-E). As these dyes provide a measure of cell permeability as well as staining extracellular DNA (TOTO-1), consistent with previous studies (17), we interpret these results to indicate that cells in the interior were intact whereas those on the periphery had compromised membranes. In comparison, biofilms treated under PYO-reductive conditions were made up of single cell layers with no secondary structures and a greater proportion of membrane-permeable cells (Fig. 1D-E). This pattern held true for all samples.

### Reduced PYO acts synergistically with antibiotics to kill cells

Sub-MICs of a range of antibiotics were added to anoxic survival reactors. Most notably, the addition of 4μg/ml of gentamicin to PYO-reductive conditions almost fully eradicated wild type biofilms (CFU/cm^2^ ± SE, n = 3, for OC = (1.71 ± 0.59) x 10^4^, - 400 mV = 21.4 ± 10.7), but did not affect Δ*phz\** biofilms. As PYO has been shown to confer tolerance to aminoglycosides (18), our data suggests that PYO confers tolerance to aminoglycosides only when it is able to cycle between reduced and oxidized states, which is also disabled when PYO is biochemically altered (14). Alternatively, or in addition, the toxicity of reduced PYO plus exposure to aminoglycosides enhances cell death. Treatment with 1 μg/ml meropenem (a β-lactam) or 4 μg/ml ciprofloxacin (a fluoroquinolone) under PYO-reductive conditions also decreased cell survival compared to OC by ~100x (CFU/cm^2^ ± SE, n = 3, for meropenem OC = (7.72 ± 0.95) x 10^4^, −400 mV = (6.73 ± 1.47) x 10^2^; for ciprofloxacin OC = (1.39 ± 0.11) x 10^4^, −400 mV = (3.53 ± 0.66) x 10^2^) but cell clusters sparsely covering the electrode surface were still present (Fig. 2B). Consistent with our results here, previously reports have shown that PYO confers tolerance to ciprofloxacin and the β-lactam carbenicillin, with Δ*phz\** cells in either colony biofilms or liquid cultures being more susceptible to ciprofloxacin and carbenicillin than wild type (18, 19). Future studies will investigate how antibiotic treatment in the absence of phenazines compares to the conditions presented here, where PYO is present, but in a reduced state. Treatment with 10 μg/ml colistin did not show an additive effect with PYO-reductive conditions, yet, PYO-oxidative conditions in the presence of colistin showed a 10x decrease in CFUs compared to non-treated biofilms (CFU/cm^2^ ± SE, n = 3, +100 mV + 10 μg/ml colistin = (1.74 ± 0.15) x 10^4^, +100 mV no antibiotic = (1.91 ± 0.06) x 10^5^). Previous studies have shown that colistin synergizes with phenazines to kill cells in colony biofilms under oxic conditions (18). Based on our results under anoxic conditions with cell-permeability dyes, we hypothesize PYO-oxidative conditions are most likely to mimic those within colony biofilms since a metabolically heterologous community with a larger proportion of metabolically active cells arises when oxidized PYO is available (Fig. 2B) and this resembles what we would expect for cells within colony biofilms grown under oxic conditions.

**Figure 2.**
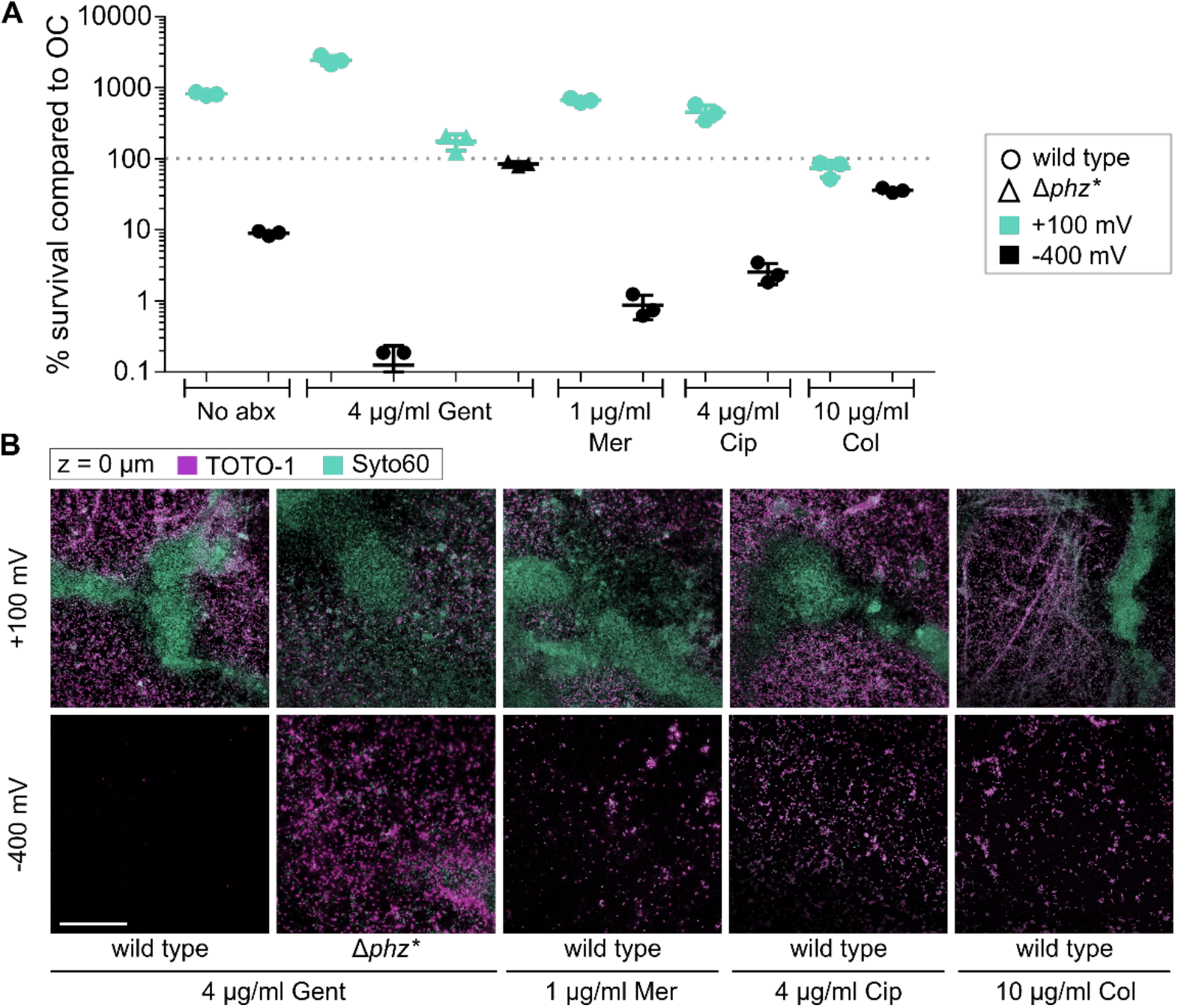
Reduced PYO acts synergistically with antibiotic treatment. **A**) CFUs after 72 hours under anoxic conditions normalized to parallel OC samples (see raw data in Fig. S3) in the presence of either 4 μg/ml gentamicin (Gent), 1 μg/ml meropenem (Mer), 4 μg/ml ciprofloxacin (Cir), or 10 μg/ml colistin (Col), n=3. Error bars represent standard error. Data from samples not treated with antibiotics (No abx) from Fig. 1 plotted again for ease of comparison. **B**) Fluorescence microscopy images of biofilm interface with electrode surface using TOTO-1 (cell-impermeable, eDNA) and Syto60 (cell-permeable, all DNA) from representative samples shown in A). Bar = 50 um.

To characterize the effect of reduced PYO on cell survival over time, biofilms were harvested pre-transfer under oxic conditions and after 30 minutes, 6, 36, and 72 hours from transfer to anoxic conditions. Plating was done in parallel on both oxic and anoxic media to rule out effects of experimental setup on cell death. Pre-transfer PYO-reductive conditions showed no significant change compared to PYO-oxidative conditions. After 30 minutes from transferring to anoxic reactors, CFUs from biofilms grown under PYO-reductive conditions decreased by 100x compared to original aerobic biofilms, while PYO-oxidative conditions only caused a slight decrease in CFUs (Fig. S4). These results indicate that under anoxic conditions, reduced PYO might be toxic, therefore, we used mid-log aerobic cell cultures to inoculate anoxic medium containing a biochemical O2 scavenging system and increasing concentrations of reduced PYO; this treatment led to a 2-3x decrease in CFUs (Fig. S5). However, increasing concentrations of reduced PYO did not correlate with a decrease in CFUs and did not achieve the 10x decrease seen in biofilm experiments between OC and PYO-reductive conditions. The discrepancy between liquid culture and biofilm experiments may be due to 1) an anti-toxicity pathway present in fresh liquid cultures but unexpressed in cells within week-old biofilms or 2) the presence of a working electrode constantly driving the PYO pool towards a fully reduced state or creating secondary toxic products at the biofilm attachment surface.

Our results highlight the importance of EET for *P. aeruginosa* survival within oxygen-limited biofilms and demonstrate that electrochemical manipulation, in tandem with antibiotic treatment, can be applied to better control biofilms of opportunistic pathogens. This work also provides context for the mechanism behind previous observations of cell death in the presence of a weak electric current (10) and provides conditions under which existing electrical bandage technology (11) may be modified to become more host-compatible. Finally, several novel research queries are also posed by the data presented here, such as characterizing the mechanism of toxicity of reduced PYO and how it synergizes or antagonizes the effect of particular antibiotics as well as studying the effect of electrochemically blocking electron shuttle cycling of borrowed phenazines to non-Pseudomonads.

## Acknowledgements

We thank Georgia Squyres, Richard Horak, and Jinyang Li for helpful feedback on experimental design and an earlier version of the manuscript. Support for this project was provided by a grant from the NIH to DKN (1R01AI127850-01A1).

